# The influence of cyclic tensile strain on multi-compartment collagen-GAG scaffolds for tendon-bone junction regeneration

**DOI:** 10.1101/406959

**Authors:** William K. Grier, Raul A. Sun Han Chang, Matthew D. Ramsey, Brendan A.C. Harley

## Abstract

Orthopedic injuries often occur at the interface between soft tissues and bone. The tendon-bone junction (TBJ) is a classic example of such an interface. Current clinical strategies for TBJ injuries prioritize mechanical reattachment over regeneration of the native interface, resulting in poor outcomes. The need to promote regenerative healing of spatially-graded tissues inspires our effort to develop new tissue engineering technologies that replicate features of the spatially-graded extracellular matrix and strain profiles across the native TBJ. We recently described a biphasic collagen-glycosaminoglycan (CG) scaffold containing distinct compartment with divergent mineral content and structural alignment (isotropic vs. anisotropic) linked by a continuous interface zone to mimic structural and compositional features of the native TBJ. Here, we report application of physiologically relevant levels of cyclic tensile strain (CTS) to the scaffold via a bioreactor leads to non-uniform strain profiles across the spatially-graded scaffold. Further, combinations of CTS and matrix structural features promote rapid, spatially-distinct differentiation profiles of human bone marrow-derived mesenchymal stem cells (MSCs) down multiple osteotendinous lineages. CTS preferentially upregulates MSC activity and tenogenic differentiation in the anisotropic region of the scaffold. Further, there are no negative effects of CTS on MSC osteogenic potential in the mineralized region previously shown to promote robust bone regeneration. Together, this work demonstrates a tissue engineering approach that couples instructive biomaterials with physiological stimuli as a mean to promote regenerative healing of orthopedic interfaces.

## 1. Introduction

The tendon-bone junction (TBJ), or enthesis, is a complex stratified region that functionally integrates tendon to bone to provide a smooth transition between the two dissimilar tissues ^1, 2^. The aligned and elastic structure of tendon makes it strong under tensional loading, but it inserts into mineralized bone that exhibits a multiple order of magnitude increased moduli. As a result, the interface between these two tissues is susceptible to high stress concentrations ^3^. In order to effectively dissipate these high stress concentrations while maintaining structural integrity, the TBJ is a compliant transitional tissue that contains gradients in collagen alignment and local mineral content^4, 5^. Nevertheless, certain TBJs, such as in the rotator cuff, are prone to a variety of chronic and acute injuries. Unfortunately, the intricate nature of the enthesis is not naturally regenerated following surgical repair, which is characterized by the formation of disorganized scar tissue, resulting in high rates of recurrence ^5, 6^. Improved techniques for the regeneration of the full spectrum of tendon-to-bone need to be developed to facilitate future studies to promote regeneration of a functional osteotendinous interface.

Tissue engineering methods provide an attractive strategy to improve current standards for tissue interface regeneration associated with spatially-graded tissues. The extracellular matrix (ECM) environment in which a cell resides has been shown to be critical for tissue development by influencing cell proliferation, morphology, adhesion, and differentiation ^7, 8^. As a result, recent efforts associated with tendon-bone repair have concentrated on efforts to develop polymeric biomaterial interfaces containing gradients of mineral content ^9^ or multiphase materials comprised of distinct materials optimized for tendon, bone, and interface regeneration ^10, 11^. Efforts in our laboratory have been inspired by collagen-GAG (CG) scaffolds which can be fabricated by freeze-drying acidic suspensions of type I collagen and glycosaminoglycans ^12^. Composed of natural ECM components, CG scaffolds have shown great promise as tissue engineering scaffolds for dermal ^12^ and nerve ^13^ regeneration as well as applications in cartilage repair ^14^. Recently, efforts in our lab have developed a directional solidification approach to generate CG scaffolds containing aligned tracks of ellipsoidal pores that replicate features of the anisotropic tendon ECM, promote transcriptomic stability of tenocytes, and which promote tenogenic differentiation of mesenchymal stem cells (MSCs) in the absence of growth factor supplements ^15-19^. We also have reported a calcium phosphate-mineralized (CGCaP) scaffold variant that have been shown to robustly promote MSC osteogenic differentiation and bone regeneration in the absence of traditional osteogenic supplements (e.g., BMP-2) ^20-25^. Combining directional solidification with a layering approach previously developed to fabricate spatially-graded CG scaffolds with mineralized and non-mineralized regions for osteochondral repair ^26, 27^, we recently reported a multi-compartment CG scaffold for TBJ applications that contains distinct regions of structural anisotropy and mineral content linked by a continuous, graded interface ^28^. This scaffold promotes initial stages of spatially-selective tenogenic and osteogenic differentiation of human mesenchymal stem cells (MSCs), indicating that it would be a promising candidate for future studies to regenerate the full spectrum from tendon to bone.

Strategies to promote robust, multi-lineage differentiation of MSCs down osseous and tendinous lineages in a spatially controlled manner must also consider clinical conditions and limitations. Generally, soluble growth factors are used to induce a strong tenogenic (e.g., GDF-5/7) ^17, 29, 30^ or osteogenic (e.g., BMP-2) ^31, 32^ response. Growth factors present complications for clinical applications due to their prohibitive cost, dosage requirements, and off-target effects ^33, 34^. Mechanical forces are essential during the development of the TBJ ^35, 36^, as well as to promote rehabilitative healing after injury ^37, 38^. As a result, efforts are increasingly exploring the application of low-amplitude cyclic tensile strain (CTS) in order to reproduce the unique mechanical environment of tendons and promote more efficient cell activity ^39-42^. Mechanical strain can promote phosphorylation of ERK 1/2 mitogen-activated protein kinase (MAPK) pathway via the activation of RhoA ^43^, leading to the upregulation of procollagen mRNA ^44^. A number of studies have focused on identifying strain paradigms to maximize ERK 1/2 activation^39^, often using intermittent strain paradigms to reduce p38 MAPK related inhibition of ERK 1/2 which can occur under extended exposure to strain ^44, 45^. Notably, *Paxton et al.* demonstrated that tendon fibroblasts seeded in fibrin hydrogels showed increased ERK 1/2 activation when exposed to an intermittent cyclic tensile strain paradigm (10% strain at 1 Hz for just 10 minutes every 6 hours) for ligament engineering applications ^39^. We recently reported the development of a CTS bioreactor to apply the same CTS profile (10% strain, 1 Hz, 10 minutes every 6 hours) to promote MSC tenogenic differentiation in the anisotropic CG scaffold^18^.

Here, we integrate a spatially-graded CG biomaterial with a CTS bioreactor system to examine synergies between the application of an intermittent strain paradigm and a non-uniform scaffold environment on TBJ-associated MSC differentiation. Here, discrepancy in the elastic moduli between the anisotropic (*tendinous*) and mineralized (*osseous*) compartments suggests the application of bulk CTS will lead to different local strain profiles across the scaffold. While we have previously described long-term culture of individual tendinous^17, 18^ and osseous^23^ scaffold regions as well as MSC activity in an osteotendinous scaffold where constant strain was applied across the entire scaffold^46^, here we focus on examining early stage MSC differentiation patterns, via signal transduction and gene expression analyses, across the spatially-graded osteotendinous scaffold when exposed to compartment specific CTS profiles. We report the combined effects of CTS and scaffold microstructure on spatially-selective MSC proliferation, mechanotransduction pathway activation, and osteo-tendinous differentiation patterns over short-term (<7 days) *in vitro* culture. We further examine preliminary development of an enthesis phenotype at the interface between the two compartments in response to CTS.

## 2. Materials and Methods

### 2.1 Preparation of collagen suspensions

Two types of collagen suspensions were prepared as previously described^46^. The first, a nonmineralized collagen-GAG (CG) suspension consisted of 1 w/v% type 1 microfibrillar collagen from bovine tendon (Collagen Matrix Inc. Oakland, NJ) and heparin from porcine intestinal mucosa (Sigma-Aldrich, St. Louis, MO), homogenized in 0.05M acetic acid at a ratio of collagen to glycosaminoglycan at 11.25:1 ^12, 47^. A collagen-glycosaminoglycan suspension containing calcium phosphate (CGCaP) was made by adding calcium salts (Ca(OH)_2_, Ca(NO_3_)_2_·4H_2_O) with the collagen and heparin and by substituting phosphoric acid as the solvent ^20, 48^. Both suspensions were homogenized at 12,000 rpm and 4°C to prevent collagen denaturation as previously described ^12, 49^. Following homogenization, both suspensions were stored overnight at 4°C and degassed prior to use.

### 2.2 Multi-compartment scaffold fabrication

Multi-compartment scaffolds were fabricated by combining a previously described directional solidification approach^15, 49^ with a liquid-phase co-synthesis method ^26^. First, degassed CG suspension was pipetted into individual wells (6mm diameter, 30mm deep) within a polytetrafluoroethylene (PTFE) mold with a copper bottom. The CGCaP suspension was subsequently layered on top, and the mold placed on a freeze-dryer shelf (VirTis, Gardiner, NY) precooled to −10°C and maintained at this temperature for 2 h to complete freezing^15^. Following freezing, the frozen suspensions were then sublimated at 0°C and 200mTorr to remove the ice crystals, resulting in a dry porous scaffold with a distinct mineralized CGCaP and aligned CG compartments with a continuous interfacial region. Following lyophilization, a guide was used to trim the ends of each scaffold equidistant from the interface to a total length of 25 mm. The scaffolds were then stored in a desiccator until use.

Hollow end blocks were fabricated from acetyl-butyl-styrene (ABS) using a MakerBot Replicator 2X 3D printer (MakerBot Industries, Brooklyn, NY)^18^, were filled with a 5:1 solution of polydimethylsiloxane (PDMS): catalyst (Hisco Inc. Houston, TX) which was allowed to partially cure for 45 minutes at 37°C before 5 mm of the CGCaP end of the scaffold was inserted into the PDMS filled end block. The PDMS was allowed to cure overnight at 37°C with the process repeated to insert the non-mineralized CG side of the scaffold in a second endblock, leaving a 15 mm gauge length specimen. A group of 6 scaffolds was set aside and marked with black India Ink (Dick Blick Art Materials, Galesburg, IL) for analysis of strain profiles across the scaffold under CTS.

### 2.3 Scaffold hydration and crosslinking

Scaffold-end block constructs were hydrated using previously described two-step process^17^. The scaffolds were soaked in 200-proof ethanol followed by phosphate-buffered saline (PBS) overnight. After hydration, the scaffolds underwent carbodiimide crosslinking in a solution of 1-ethyl-3-[3-dimethylaminopropyl] carbodiimide hydrochloride (EDC) and N-hydroxysulfosuccinimide (NHS) at a molar ratio of 5:2:1 EDC:NHS:COOH in PBS for 2 hours at room temperature ^50, 51^. Prior to seeding, scaffolds were soaked in complete MSC growth media, consisting of low-glucose Dulbecco’s Modified Eagle Medium with 10 vol% MSC fetal bovine serum and 1 vol% antibiotic-antimitotic (Thermo Fisher, Waltham, MA), for 72 hours^18^.

### 2.4 Critical point drying and SEM analysis

Scaffolds were dehydrated in a series of ethanol washes (10% to 100%) followed by to critical point drying (CPD) using a Samdri-PVT-3D (Tousimis, Rockville, MD)^52^. Dry samples were then sectioned and mounted on carbon tape for sputter coating with a gold/palladium mixture. Imaging was then carried out with a Philips XL30 ESEM-FEG (FEI Company, Hillsboro, OR) at 5 kV with a secondary electron detector.

### 2.5 Cyclic tensile strain bioreactor

The cyclic tensile strain bioreactor used in this study was previously described for use with monolithic anisotropic CG scaffolds to identify CTS paradigms that enhanced MSC tenogenic differentiation^18^. While 10% cyclic strain (at 1 Hz for 10 minutes every 6 hours) was shown to promote tenogenic differentiation in a monolithic tendinous CG scaffold, this study applied 5% system (bulk) strain (at 1 Hz for 10 minutes every 6 hours) to the multi-compartment scaffold. This change took into account the significant difference in elastic moduli of the tendinous and osseous scaffold compartments (E_tendinous_ << E_osseous_) that resulted in increased strain concentrated in the non-mineralized tendinous compartment^46, 53^ as well as the 50:50 ratio of tendinous:osseous scaffold compartments in the composite used in this study.

The CTS bioreactor contains 24 individual wells with loading posts and a rake system connected to a programmable linear actuator controlled via a custom C# program (Pololu Corp. Las Vegas, NV)^40^. A second set of wells with fixed posts was used as static control (0% strain condition). Local scaffold strain profiles across the multi-compartment scaffolds were measured as previously described^18^. Briefly, scaffolds were marked with India Ink prior to hydration. Video of scaffold deformation was acquired using a Canon EOS-5D Mark II SLR 21.1MP Digital Camera (Canon, Tokyo, Japan) while the scaffolds were cyclically loaded to 5% system strain at 1 Hz. The distance between each mark was tracked between frames using the Object Tracker plugin in ImageJ, with strain calculated using distances between each point at rest ^18^. Groups of points within each compartment were analyzed in order to determine compartment specific strain.

### 2.6 Human mesenchymal stem cell expansion and culture in multi-compartment scaffolds under CTS

Human bone marrow-derived MSCs were acquired from Lonza (Walkersville, MD). Multiple lots originating from separate donors were pooled in order to provide sufficient numbers of cells for all experiments and to reduce the potential for any donor-specific responses. MSCs were cultured in complete MSC growth medium, at 37°C and 5% CO2, fed twice a week, and used at passage 6 for all experiments.

Scaffold-end block constructs (6 mm diameter, 15 mm gauge length, 50:50 tendinous-osseous compartment) were seeded with MSCs using a previously validated static seeding method ^54^. Briefly, hydrated scaffolds were gently blotted to absorb excess liquid and then seeded with 3.0×10^5^ MSCs per 60 μL media (3×20 μL drops along scaffold length) in the individual wells of the bioreactor and static well systems. Cells were then incubated at 37°C and allowed to attach for 2 hours before the addition of media to the wells; constructs were then given an additional 24 hours before initial application of CTS to ensure cell attachment. Cell-seeded scaffolds in the CTS groups were subjected to continuous cyclic tensile strain (5% overall strain at 1 Hz) for 10 minutes every 6 hours, for up to 6 days, using a previously described protocol^18^ inspired by *Paxton et al.* ^39^. MSCs were maintained in osteotendinous scaffolds in the bioreactor (or static control well) for up to 6 days at 37°C and 5% CO_2_ in complete MSC growth media (replaced 2x/week) that was not supplemented with either osteogenic or tenogenic growth factors. During the experiment, scaffolds were collected for further analysis at designated times, or immediately after the conclusion of a strain cycle. Osteotendinous scaffolds were collected from the bioreactor at timepoints up to 6 days. A cutting guide was used to consistently cut the scaffolds into three equally sized sections for compartment-specific analyses with a razor blade: non-mineralized CG (*tendinous*), mineralized CGCaP (*osseous*), and the middle third of the scaffold containing the interface as well as adjacent CG and CGCaP regions (*interface*). While the gradient transition between *osseous* and *tendinous* scaffold compartments in our osteotendinous scaffold is on the order 250µm ^46^, to extract sufficient RNA and protein for analysis the *interface* zone analyzed here is comprised of that interface along with adjacent regions of aligned and mineralized scaffold. As a result, for this work we primarily focus on examining divergent responses of MSCs in the *tendinous* vs. *osseous* compartments.

#### 2.6.1. Quantifying metabolic activity of MSC-seeded scaffold sections

The mitochondrial metabolic activity of the MSCs within each scaffold section was measured via non-destructive alamarBlue® assay ^55^. Cell-seeded sections were incubated in 10% alamarBlue (Invitrogen, Carlsbad, CA) solution with gentle shaking for 1 hour. Metabolic activity was compared to a standard curve generated from known cell numbers at the time of seeding and reported as a percentage of the total number of seeded cells.

#### 2.6.2. Protein extraction, gel electrophoresis, and immunoblotting

Cellular proteins were extracted from each scaffold region using a previously described protocol^18, 46^. Samples were briefly washed in warm PBS, blotted dry, and then submerged in RIPA buffer containing protease and phosphatase II and III inhibitor cocktails. For SDS PAGE, 1:1 solutions of Laemmli sample buffer and 10 µg of protein in RIPA buffer were heated to 90°C for 10 minutes then loaded onto 4-20% gradient tris-glycine and separated using a constant 150V for 90 minutes. A semi-dry transfer was used to transfer proteins to a nitrocellulose membrane (GE Healthcare) at 15 V for 15 minutes. Membranes were then blocked for 1 hour in 5% milk in Tris-buffered-saline + 0.1% Tween (TBST), incubated overnight at 4°C with the appropriate primary antibody in 5% bovine serum albumin in TBST, washed in TBST, then incubated for 1 hour at room temperature with an HRP-linked goat anti-rabbit IgG secondary antibody (1:5,000. Cell Signaling) in TBST. Antibody binding was then detected using SuperSignal West Pico or Femto Chemiluminescent Substrates (Thermo Fisher, Waltham, MA) on an Imagequant LAS 4010 system (GE Healthcare). Primary and secondary antibodies were stripped with OneMinute® Western Blot Stripping Buffer (GM Biosciences, Rockville, MD) so membranes could be re-probed up to two times, following the same protocol. Antibodies were purchased from Cell Signaling (Danvers, MA): p38 (8690), p-p38 (9215), ERK 1/2 (4695), pERK 1/2 (4370), Smad 2/3 (8685), pSmad 2/3 (8828), Smad 1 (6944), pSmad 1/5/8 (9511), β-actin (4967; control). ImageJ was used to quantify intensity of bands.

#### 2.6.3. RNA isolation and real-time PCR

RNA was extracted from scaffold sections on days 1, 3, and 6 using an RNeasy Plant Mini kit (Qiagen, Valencia, CA). The mRNA was then reverse transcribed to cDNA using the QuantiTect Reverse Transcription kit (Qiagen) in a Bio-Rad S1000 thermal cycler as previously described^16, 56^. Real-time PCR reactions were carried out in triplicate (10ng of cDNA) using the QuantiTect SYBR Green PCR kit (Qiagen) in a 7900HT Fast Real-Time PCR system (Applied Biosystems, Carlsbad, CA). The primers used were consistent with previous studies, and were synthesized by Integrated DNA Technologies (Coralville, IA). The expression level of the following markers was quantified: collagen type III alpha 1 (*COL3A1*), cartilage oligomeric matrix protein (*COMP*), scleraxis (*SCXB*), Mohawk homeobox (*MKX*), aggrecan (ACAN), SRY Box 9 (SOX9), alkaline phosphatase (*ALP*), bone sialoprotein (*BSP*), osteopontin (*OP*), runt-related transcription factor 2 (*RUNX2*), and glyceraldehyde 3-phosphate dehydrogenase (*GAPDH*), which was used as a house keeping gene (**Table 1**). Results were generated using the ΔΔCt method, and all results were expressed as fold changes normalized to the expression levels of MSCs at the time of seeding the scaffolds.

**Table 1:**
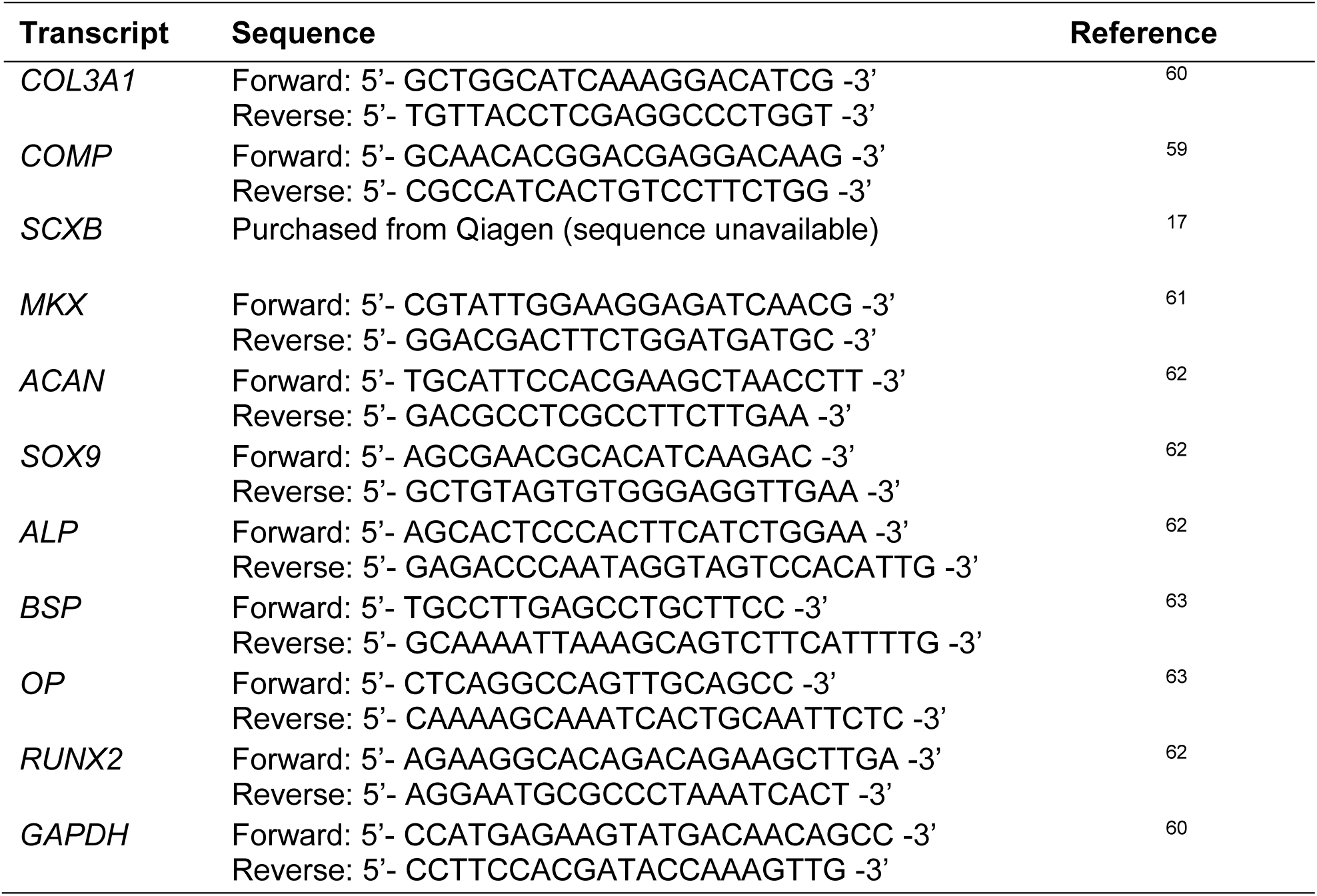
PCR primer sequences.

### 2.7 Statistics

Statistical analysis of parametric data was performed using two-way analysis of variance (ANOVA) on the western blot, metabolic activity, and gene expression data sets to evaluate the effects of scaffold compartment and strain conditions (independent variables: scaffold compartment, strain condition). One-way ANOVA was performed within each group to evaluate temporal effects (independent variable: time). ANOVA was followed by Tukey-honest significant difference *post hoc* tests. Significance was set at *p* < 0.05. A minimum of three independent scaffolds were analyzed at each time point for all metrics. Error is reported in figures as the standard error of the mean. All statistical analysis was performed in Microsoft Excel using the Real Statistics Resource Pack.

## 3. Results

### 3.1 Application of cyclic tensile strain across spatially-graded scaffolds

Multi-compartment scaffolds, fabricated using a method previously described by our group^28^, contain distinct regions of structural anisotropy and mineral content as well as a continuous interface **(Figure 1)**. Scaffolds were embedded into endblocks prior to loading into a custom cyclic tensile strain bioreactor with individual wells for each scaffold that would be attached to a sliding rake system controlled by a linear actuator^18^ **(Figure 2A)**. As a control, scaffolds were alternatively placed into identical static well plates that remained unloaded throughout **(Figure 2B)**. Due to the spatially-graded differences in the stiffness of the osteotendinous scaffold, we quantified strain profiles across the entire scaffold as well as local strain profiles in the tendinous (aligned, non-mineralized) and osseous (mineralized) scaffold compartments. A system setting of 5% strain translated to a 3.44 ± 0.05% overall (bulk) strain with 5.69 ± 0.04% and 1.90 ± 0.11% strain in the CG and CGCaP compartments, respectively. The discrepancy in system setting (movement of the sliding rake: 5%) and the bulk strain on the scaffold (3.44%) was due to tolerances required to attach the scaffolds to the rake system ^18^. Throughout the remainder of the manuscript, we will refer to the magnitude of applied strain as the movement of the sliding rake system (system setting).

**Figure 1.**
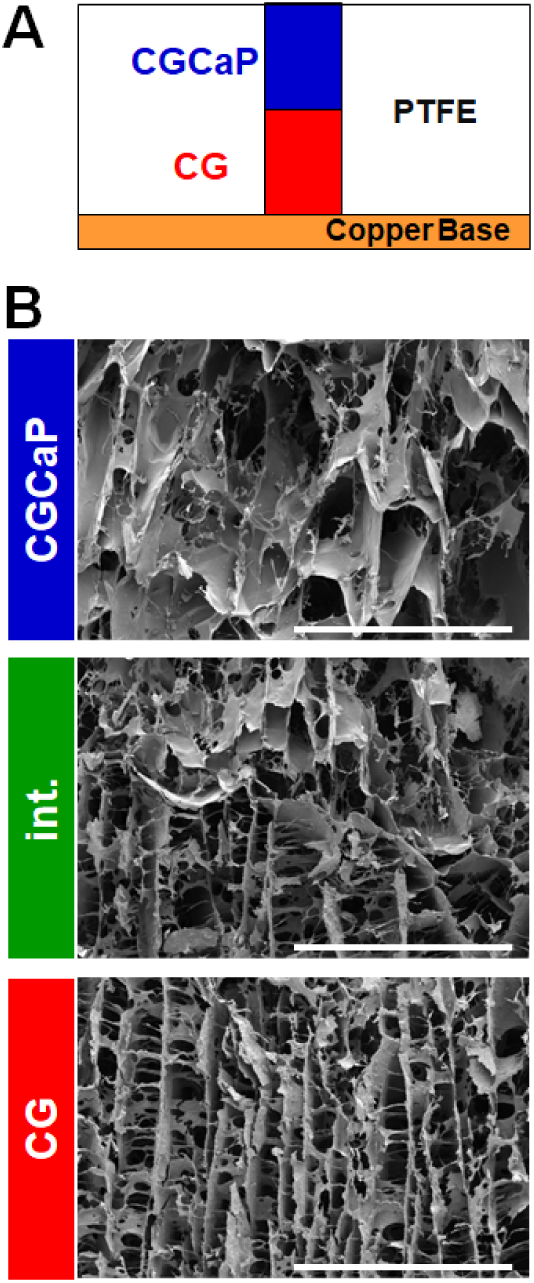
**(A)** Schematic of scaffold mold with layered collagen suspensions. **(B)** Scanning electron micrographs of each zone of a multi-compartment CG scaffold (tendinous CG; interface; osseous CGCaP). Scale bars = 1 mm.

**Figure 2.**
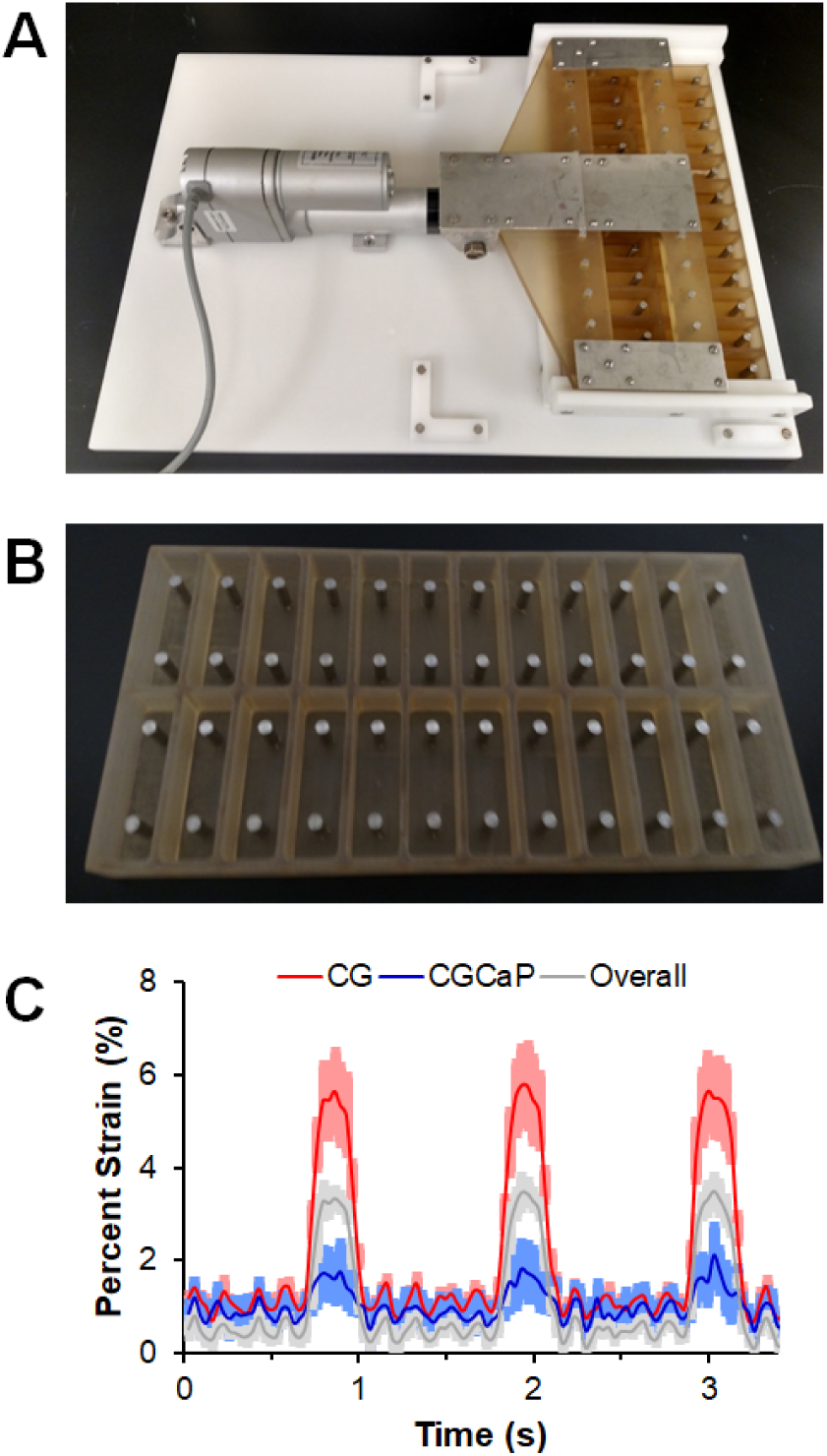
**(A)** Custom bioreactor designed with 24 individual wells with static loading posts and moveable rake controlled by a programmable linear actuator. **(B)** Static well system for control cultures run in parallel to bioreactor. **(C)** Compartment-specific scaffold deformation in response to cyclic tensile strain (5%, at 1 Hz). Scaffold deformation (sample size: n=3) is shown as mean ± SEM.

### 3.2 Effects of intermittent cyclic tensile strain on compartment-specific cellular metabolic activity

MSC metabolic activity was monitored in each scaffold compartment over the course of 6 days of culture with and without CTS (5% total strain, 1 Hz, 10 minutes every 6 hours; **Figure 3)**. In general, higher levels of cellular metabolic activity were observed in the CG compartment (*tendinous*) compared to the CGCaP compartment (*osseous*). Further, MSC metabolic activity within the CG compartment was higher with CTS compared to static culture. As early as day 1 of culture, MSCs in the CG compartment subjected to CTS show significantly higher metabolic activity (p < 0.05) compared to all other groups. By day 3, cells in the static CG show higher activity compared to day 1 (p < 0.05), and cells in both CG compartments, with and without CTS, are significantly more active than in the CGCaP compartments (p < 0.05). At the end of the culture on day 6, cells in the static CG compartment and CGCaP compartment with CTS display an increased metabolic activity in comparison to day 1 (p < 0.05). MSCs in the CG compartment show increased metabolic activity with CTS compared to both days 1 and 3 as well as all other groups on day 6 (p < 0.05).

**Figure 3.**
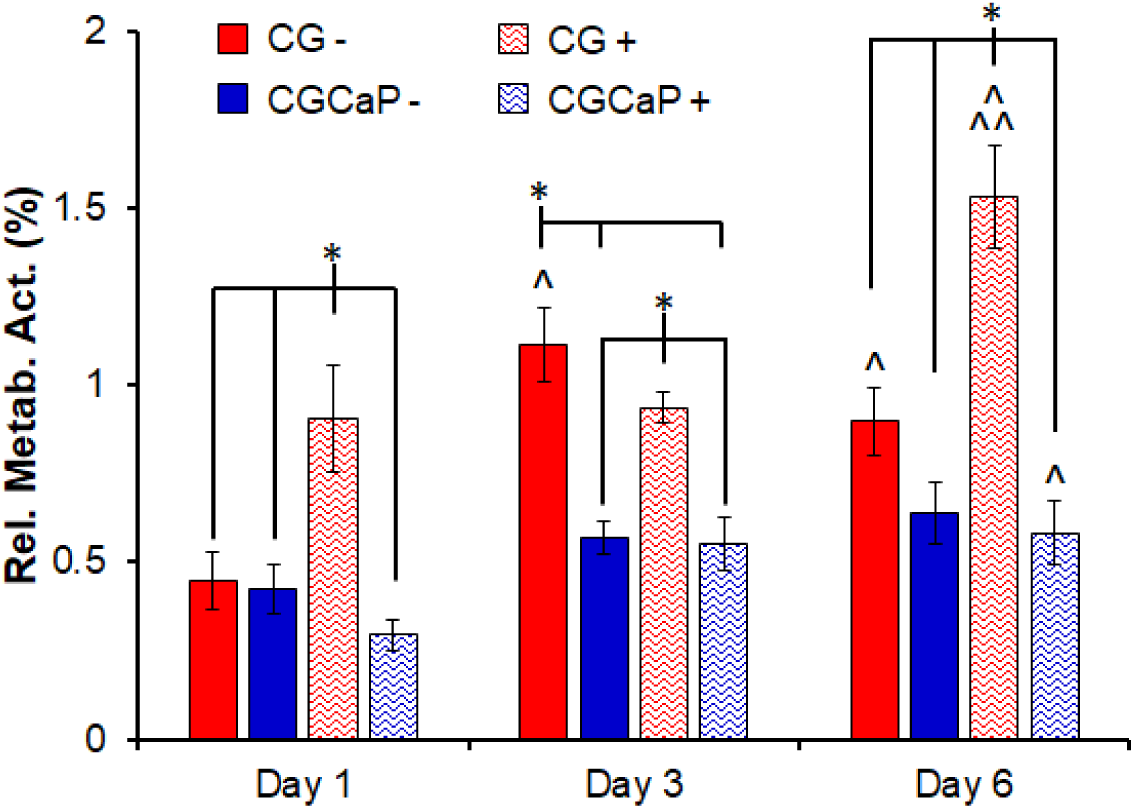
Compartment-specific metabolic activity of MSCs across the osteotendinous scaffold (CG, interface, CGCaP) in either static culture or exposed to CTS (10min at 5% strain and 1 Hz) followed by 5 hours 50 minutes recovery at days 1, 3, and 6. Data expressed as mean ± SEM (*n* = 6). ^*^ Significantly higher (*p* < 0.05) than indicated groups at same day. ^^^ Significantly higher (*p* < 0.05) than same group on day 1. ^^^^ Significantly higher (*p* < 0.05) than same group on day 3.

### 3.3 CTS promotes spatially-selective mechanotransduction and Smad pathway activation across the scaffold

We subsequently examined compartment-specific changes in mechanotransduction pathway activation (ERK1/2 vs. P38) in response to CTS. Samples were analyzed immediately prior to and at the conclusion of the first (before: 0 minutes; after: 10 minutes) and second (before: 360 minutes; after: 370 minutes) strain cycles to evaluate early activation of compartment-specific mechanotransduction pathways in response to intermittent CTS. While not significant, ERK 1/2 activation tended to be higher in the CGCaP compartment compared to the CG compartment at all timepoints (p = 0.18) **(Figure 4A)**. Additionally, immediately after the second strain cycle, ERK 1/2 activation is significantly upregulated in the CGCaP compartment (p < 0.05). The activation of p38 MAPK displays an inverse trend compared to ERK 1/2 with slightly lower activation levels in the CGCaP compartment compared to the CG compartment, and significantly reduced activation in the CGCaP compartment after the second strain cycle (p < 0.05) **(Figure 4B)**.

**Figure 4.**
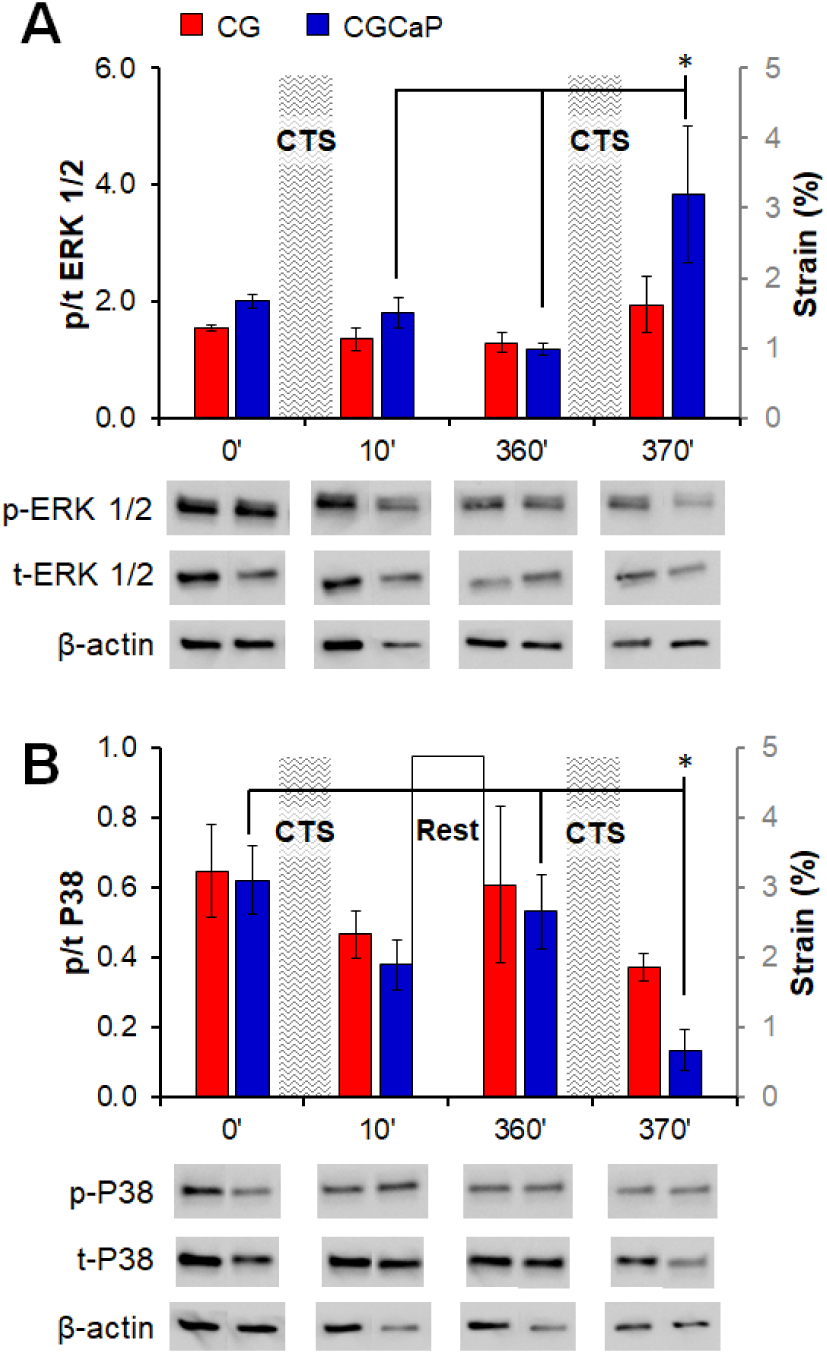
Relative levels of phosphorylated:total **(A)** ERK 1/2 and **(B)** p38 MAPK in each compartment across the osteotendinous scaffold (CG, interface, CGCaP) with and without CTS as determined by immunoblot immediately prior to and following the first and second strain cycles. Data expressed as mean ± SEM (*n* = 3). *: *p* < 0.05.

We subsequently examined compartment-specific activation of the Smad 2/3 and Smad 1/5/8 pathways after 1 and 3 days in culture, associated with increased tendinous and osseous differentiation respectively^57, 58^. While not significant, the activation of Smad 1/5/8 tended to be higher in the CGCaP compartment compared to the CG compartment and with strain compared to static condition **(Figure 5A)**. Activation of Smad 2/3 increased in the *tendinous* CG scaffold compartment with CTS after 3 days **(Figure 5B)**.

**Figure 5.**
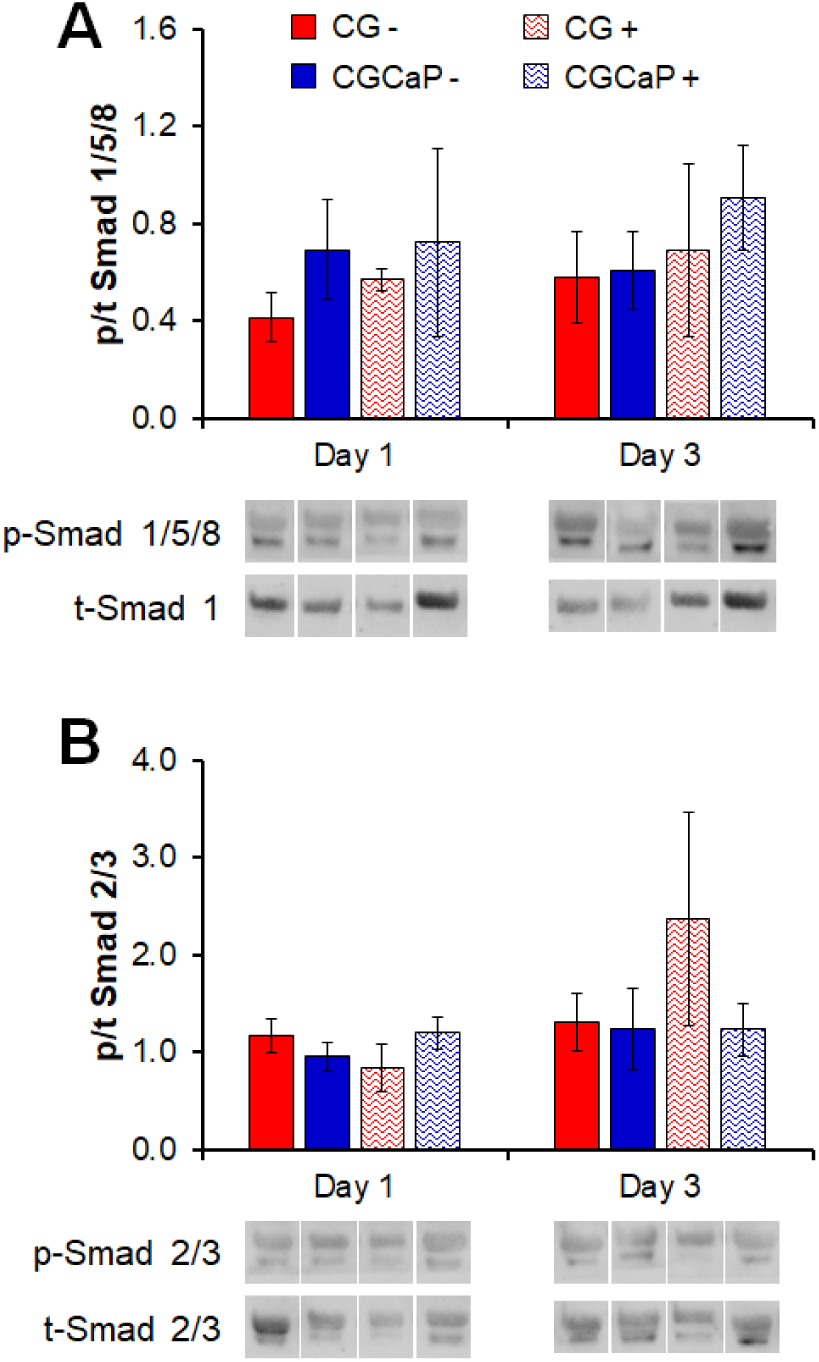
Relative levels of phosphorylated:total **(A)** Smad 1/5/8 and **(B)** Smad 2/3 in each compartment of the osteotendinous scaffold with and without CTS as determined by immunoblot on days 1 and 3. Data expressed as mean ± SEM (*n* = 3). *: *p* < .05.

### 3.4. CTS promotes compartment-specific gene expression patterns

We examined time dependent changes in gene expression profiles for MSCs in each scaffold compartment throughout the 6-day culture using a panel of tendon (*COL3A1, COMP, SCXB, MKX*, **Figure 6**)^17, 59-61^, osteogenic (*ALP, BSP, OPN, RUNX2*, **Figure 7**)^17, 62^, and fibrocartilage (*ACAN, SOX9*, **Figure 8**)^62, 63^ genes relevant to TBJ applications.

**Figure 6.**
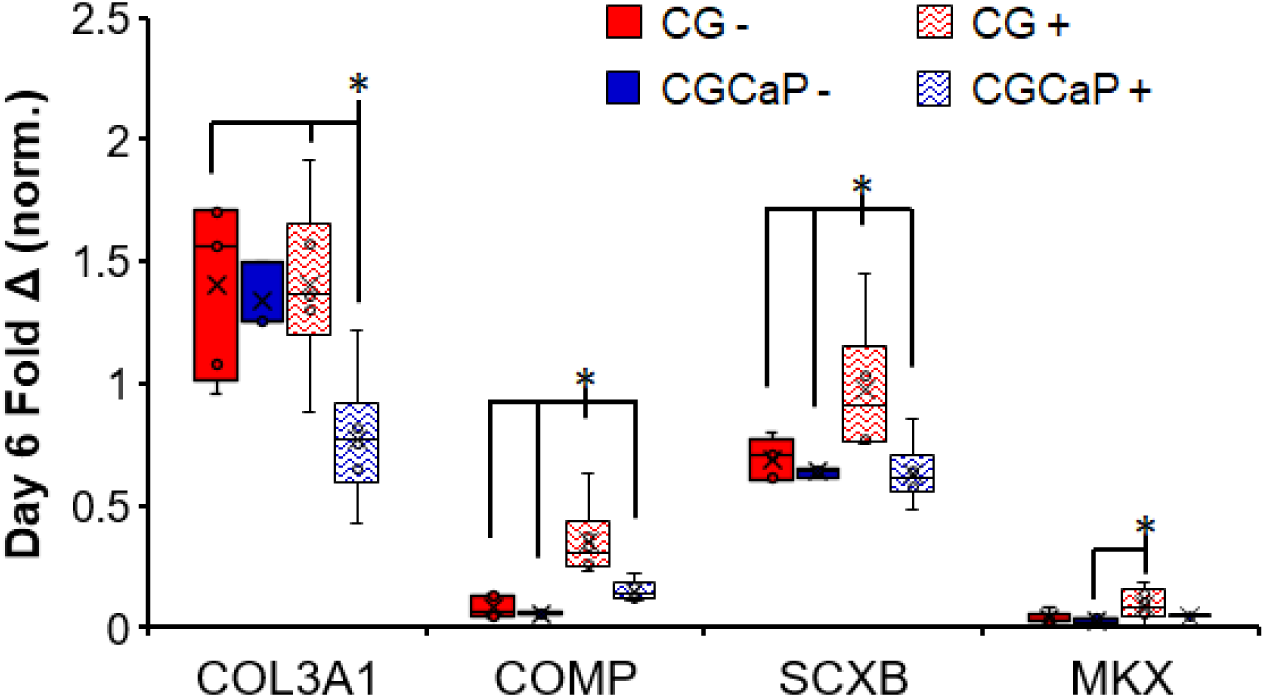
Compartment-specific expression levels of tenogenic-associated genes *COL1A1, COMP, SCXB*, and *MKX* across the osteotendinous scaffold with and without CTS on day 6. Data expressed as mean ± SEM (*n* = 6). *: *p* < 0.05.

**Figure 7.**
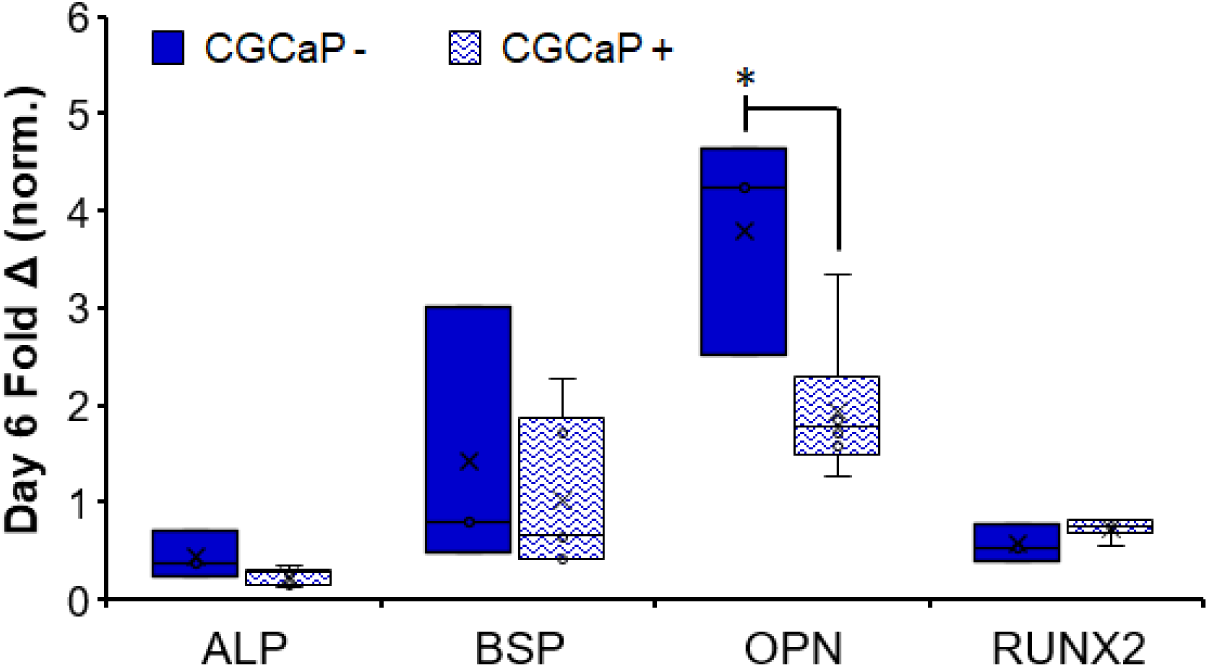
Compartment-specific expression levels of osteogenic-associated genes *ALP, BSP, OPN*, and *RUNX2* in the CGCaP (osseous) scaffold zone with and without CTS on day 6. Data expressed as mean ± SEM (*n* = 6). *: *p* < 0.05.

**Figure 8.**
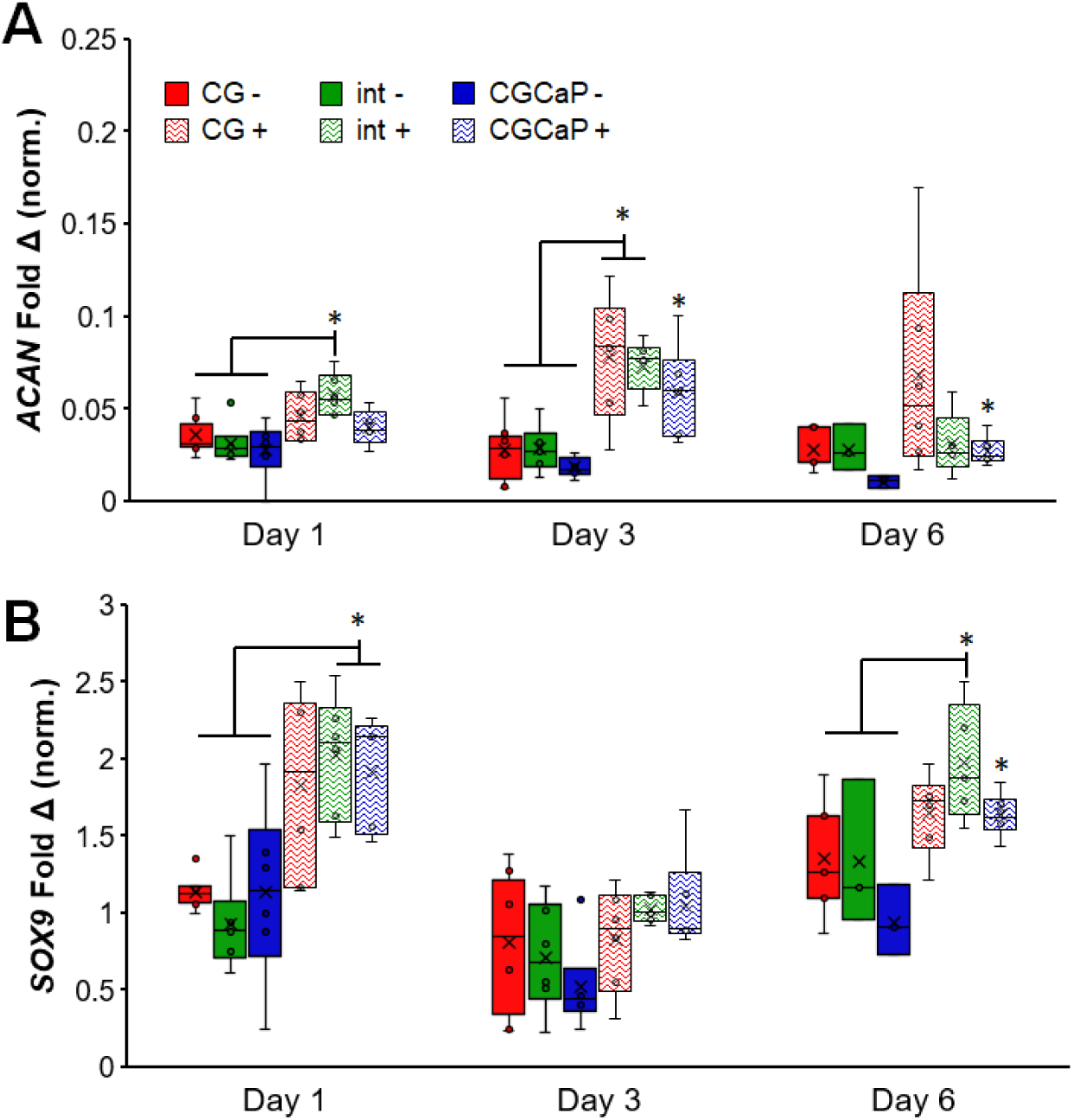
Compartment-specific expression levels of fibrocartilage-associated genes **(A)** *ACAN* and **(B)** *SOX9* across the osteotendinous scaffold with and without CTS on days 1,3, and 6. Data expressed as mean ± SEM (*n* = 6). *: *p* < 0.05.

#### 3.4.1 Tenogenic gene expression enhanced in tendinous scaffold compartment

After 1 day of culture, gene expression levels of *COMP, SCXB*, and *MKX* were generally highest in the CG and interface compartments with strain while there was negligible impact of strain on the expression of *COL1A1* **(Figure S1)**. By day 6 of CTS, we observed more marked changes in tenogenic gene expression profiles with and without CTS (**Figure 6**). A significant decrease in *COL1A1* expression is observed in the CGCaP compartment when exposed to CTS, compared to the CG compartment with or without CTS (*p* < 0.05). The expression levels for both *COMP* and *SCXB* are significantly upregulated in the CG compartment with CTS, compared to the same compartment under static conditions, and the CGCaP compartment regardless of strain conditions (*p* < 0.05). Finally, *MKX* expression, while still relatively low, is significantly higher in the CG compartment with CTS compared to the CGCaP compartment under static conditions (*p* < 0.05). Together, these results show higher expression of tenogenic differentiation associated genes in the CG (*tendinous*) scaffold compartment in response to CTS (vs. static CG), but also significantly increased expression of tenogenic associated genes in the *tendinous* vs. the CGCaP (*osseous*) compartments under CTS.

#### 3.4.2 Osteogenic gene expression maintained in osseous scaffold compartment

The CGCaP scaffold on its own has been previously shown to robustly promote MSC osteogenic differentiation ^64, 65^, so here we confirmed CTS does not significantly reduce osteogenic differentiation potential. CTS had a much smaller impact on osteogenic genes compared to the tenogenic markers over the full course of the experiment (**Figure S2**). There was no significant impact of CTS, nor scaffold compartment, on the expression levels of *BSP*. Expression levels for *OP* were higher in the CGCaP compartment with CTS, on day 1, compared to all groups under static conditions, while the interface region in static culture showed significantly lower expression than all other groups and conditions (*p* < 0.05). On days 3 and 6, this trend reversed, and the groups exposed to CTS showed lower expression levels of *OP* compared to those in static culture (*p* < 0.05). *RUNX2* expression levels were higher in the CGCaP compartment with CTS compared to the same compartment in static culture on day 1 (*p* < 0.05) but lower in the static CGCaP compartment compared to the other groups under static conditions on day 6 (*p* < 0.05). On the final day of the study, *ALP* expression was significantly higher in the interface region under static conditions compared to the other groups (*p* < 0.05). Most importantly, we examined the differences between the CGCaP compartment, with and without CTS, at the conclusion of the study (**Figure 7**). While there is a minor downregulation of *OP* expression in the CGCaP compartment with CTS (*p* < 0.05), otherwise we observed no significant impact of CTS on osteogenic gene expression within the CGCaP compartment.

#### 3.4.3 Interface gene expression enhanced with cyclic tensile strain

We examined the expression of interface associated genes across all three scaffold zones. The expression levels of both *ACAN* and *SOX9*, while variable, were generally higher in groups exposed to CTS compared to those in static culture (**Figure 8**). All groups in static culture showed significantly lower expression of *ACAN* compared to the interfacial region on day 1 and both the interface and CG compartment on day 3 (*p* < 0.05). *SOX9* expression was also significantly higher in the interface on day 6 and both the interface and CGCaP compartment on day 1 compared to all three compartments under static conditions (*p* < 0.05).

## 4. Discussion

The application of low-amplitude cyclic tensile strain is known to be an important regulator in the development of the native tendon and enthesis, as well as for the regulation of a number of stem cell fate decisions ^36, 66-68^. Our group has recently described the development of a multi-compartment CG scaffold material system with distinct regions of structural anisotropy and mineral content for tendon-bone-junction engineering ^28^. While these previous studies showed the ability to tailor material properties in order influence stem cell fate in a spatially-dependent manner, experiments were performed using a bioreactor that provided uniform strain across the entire scaffold, not compartment specific strain profiles that result during unconfined tension experiments. Recently, we described the development of a custom bioreactor system to apply CTS to a MSC-seeded anisotropic CG scaffold, identifying a CTS profile that promotes robust activation of mechanotransduction and TGF-β growth factor pathways known to be vital in the development of native tendon as well as increased expression of tenogenic differentiation associated genes ^18^.

The primary focus on this work was to examine how the application of CTS within a multi-compartment material may impart non-uniform strain profiles across the scaffold and promote initial determinants of tenogenic and osteogenic differentiation in a spatially-dependent manner. This project therefore focused on identifying localized mechanotransduction pathway activation followed by the downstream activation of growth factor-related pathways and early stage differentiation patterns via gene expression shifts, and used time points to define the early kinetics of these responses. While the timeframe of this work (up to 6 days *in vitro*) did not allow for the demonstration of extensive matrix remodeling and *de novo* tissue development, it has allowed us to show that MSCs seeded within a spatially-graded biomaterial containing gradations in microstructural alignment, mineral content, and bulk mechanics, will display disparate responses to tensile stimulation depending on location. Specifically, MSCs within the *tendinous* CG compartment displayed tenogenic-differentiation responses consistent with our previous work in single-compartment CG scaffolds^18^. When exposed to CTS, MSCs in the anisotropic CG material exhibited higher cellular metabolic activity, generally increased activation of the Smad 2/3 pathway, and significantly increased expression levels of tenogenic markers *COMP*, scleraxis (*SCXB*), and Mohawk (*MKX*).

We also evaluated the effect of CTS on cells within the *osseous* (CGCaP) compartment of the scaffold. The CGCaP compartment is about 20-fold stiffer than the CG compartment and experiences reduced levels of local strain during mechanical testing^69^ as well as locally reduced strain during CTS. However, the very low amplitude strain that was observed could have implications on MSC responses due to deformation induced fluid flow within the scaffold. Indeed, previous work has shown that both CTS ^70^ and shear flow ^71^ could be drivers of osteogenic differentiation of MSCs. However, here we observed little-to-no impact of CTS on MSCs within the CGCaP compartment. This was not altogether surprising. As we have previously shown, MSCs seeded on CGCaP scaffolds show increased mineral deposition and matrix remodeling along with upregulation of osteogenic markers *BSP*, osteopontin (*OPN*), and osteocalcin (*OCN*) ^21, 22^. This osteogenic activity is associated with scaffold-induced activation of the Smad 1/5/8 pathway in MSCs within the CGCaP material ^23, 24^. Any potential impact of the low amplitude strain that was observed in the CGCaP compartment with CTS was likely overshadowed by the MSC osteogenic response to the material itself.

While not the primary goal of this study, the evaluation of fibrocartilaginous markers at the intersection of the two compartments does provide intriguing insight for future studies. When evaluating the expression levels of *ACAN* and *SOX9* across each scaffold, a clear trend of increased expression when exposed to CTS was observed. This trend was especially apparent for *SOX9* in the middle third of each scaffold that contained the interface between the two materials. This is promising for next generation approaches to specifically regenerate the tendon enthesis. Previous work from *Spalazzi et al*. demonstrated the use of used a tri-culture system on a tri-phasic material with osteoblasts, chondrocytes, and fibroblasts seeded on bone, interface, and tendon specific regions ^11^. *He et al.* also developed a similar tri-lineage co-culture with MSCs seeded between osteoblasts and fibroblasts on a uniform silk scaffold for the development of a partially mineralized fibrocartilaginous interfacial zone ^72^. Recently, *Liu et al.* investigated the use of decellularized tendon matrix treated with ultrasound in order to influence rabbit MSCs towards a fibrocartilaginous-like state ^73^. While these previous works do show the potential for these multipotent MSCs to develop into enthesis tissue, there has yet to be any in-depth evaluation of a full tendon-bone-junction model that incorporates a physiologically relevant and competent enthesis. In this case, the combination of CTS along with the unique material properties at the intersection of two distinct, but continuous compartments, may be sufficient to drive that initial fibrocartilaginous response, and is the subject of ongoing efforts building on the work reported here.

While demonstrating the use of a spatially-graded biomaterial to promote regionally-specific MSC differentiation patters, this study also suggests a series of future experiments with the goal of optimizing scaffold properties and CTS for TBJ regeneration. While we observed Smad 2/3 activation in the tendinous scaffold compartment after exposure to CTS, a more in-depth understanding of the interplay between CTS and compartment-specific Smad activation should be explored. There are two possible modes of Smad activation. Canonical activation could result from the endogenous production of TGF-β growth factors produced with initial MSC differentiation events. Non-canonical methods of activation could also play a role as mechanotransduction pathways like ERK 1/2 and p38 have been shown to influence Smad activity ^25^. It is also possible that the initial non-canonical activation of Smad 2/3 by mechanical stimulation could drive an autocatalytic cycle of differentiation and endogenous growth factor production resulting in more robust Smad activation and MSC differentiation as postulated by *Allen et al*.^74^. The region-specific sampling methods described here will be essential for ongoing work to examine the balance of canonical and non-canonical Smad activation across the scaffold. Further, this project suggests the need for a series of ongoing studies to evaluate long-term matrix remodeling and *de novo* tissue development of MSCs within these scaffolds in response to CTS. Finally, based on evaluation of markers of fibrocartilage differentiation, future efforts will likely need to focus on the incorporation of a functional enthesis zone at the interface between the tendinous and osseous scaffolds. Such efforts require higher resolution methods to evaluate cellular activity and gene expression at the narrow region, which is generally less than 1 mm, between the adjacent *tendinous* and *osseous* scaffold compartments ^28, 69^.

## Acknowledgements

The authors would like to acknowledge the Carl R. Woese Institute for Genomic Biology Core Facilities for assistance with real-time PCR system and immunoblotting. This research was carried out in part at the Imaging Technology Group within the Beckman Institute for Advanced Science and Technology at the University of Illinois at Urbana-Champaign. The authors would like to thank Scott Robinson and Cate Wallace for assistance with critical point drying and scanning electron microscopy. Research reported in this publication was supported by the National Institute of Arthritis and Musculoskeletal and Skin Diseases of the National Institutes of Health under Award Number R21 AR063331. Additional support was provided by the Chemical and Biomolecular Engineering Dept. and the Carl R. Woese Institute for Genomic Biology (BACH) at the University of Illinois at Urbana-Champaign.

